# Familial heterogeneity in breast cancer predisposition: a study of 22 Utah families

**DOI:** 10.1101/092841

**Authors:** Elizabeth O’Brien, Richard A. Kerber, Raymond L. White

## Abstract

The problem of “missing heritability” in genome-wide analyses of complex diseases is thought to be attributable to some combination of: rare variants of moderate to large effect, common variants of very small effect, and epigenetic, epistatic, or shared environmental effects. Rare variants do not affect large numbers of people by definition, but identified genes and pathways frequently lead to important insights into pathogenesis, and become targets of chemoprevention or therapy. Family studies remain an efficient way to identify rare variants with sizable effects on disease risk. We present a genome-wide study of breast cancer in 22 large high-risk families including 154 women diagnosed with breast cancer. Appropriate marker spacing was achieved by simulation studies of founder haplotypes to reduce the chance that linkage disequilibrium produced spurious linkage peaks. For each family, we generated 100 simulations of null linkage genome-wide to estimate the probability that individual results were due to chance. We identified a total of 12 putative susceptibility regions with per-family genome-wide probability < 0.05. These regions were located on 10 chromosomes; 10 of the 22 families showed linkage at these locations; two or more families showed linkage to 6 regions on 5 chromosomes (4q, 5q, 6p, 14q, 18p, and 18q). These results indicate that there is considerable heterogeneity among families in genomic regions and thus variants predisposing to breast cancer. Moreover, they suggest that uncommon high– or medium-risk genetic variants remain to be found, and that family designs can be an efficient way to identify them.

## Introduction

The genetic dynamics of complex traits have concerned population scientists for more than a century, but the quantity of data streaming from genomic studies in recent decades has drawn new focus to the prospect of identifying genes underlying complex phenotypes. Especially important targets for genetic characterization are human disease phenotypes that commonly plague us and frequently kill us, such as cancer.

Long before genome-wide data were available for complex trait analysis, family studies were the workhorses used to study the genetic basis of cancer because case clusters were originally observed in families. Examination of familial clusters of neoplastic disease led to the identification of the tumor suppressor role of *TP53* [1] in Li-Fraumeni syndrome, retinoblastoma, and the role of the *FANC* gene complex in Fanconi anemia (FA). Family studies of breast cancer also provided the first plausible evidence that a few genes of at least moderate effect might account for excess risk and observed case aggregation in families. This result was established for *BRCA1* and *BRCA2* mutations in familial breast and ovarian cancers [2] [3], and as a result the two genes were dubbed “most important” for breast cancer predisposition in high risk families [4].

Although breast cancer is not the most common of FA’s neoplastic effects, it has been demonstrated fairly recently that the products of the *FANC* gene complex function in congress with BRCA1 and BRCA2 in DNA repair pathways and provisionally explains their concordant effects on breast cancer predisposition in some families [5]. Mutations in *PTEN* and *STK11* may also exhibit relatively high penetrance effects [6–9] while other genes, such as *ATM*, *CHK2*, and *PALB1*, also account for excess breast cancer risk in some families with somewhat lower penetrance[10, 11]; however, families segregating these other mutations are rarer, and thus account for less of the total genetic risk estimated for large and heterogeneous case series. In fact, no other genes as commonly mutated, or of such high penetrance as *BRCA1* and *BRCA2,* have been identified yet through family studies of breast cancer. Therefore, it has generally been concluded from numerous studies of familial cancer risk (breast and other) in multiple populations, that: 1) the same genes do not account for cancer incidence in all families with elevated risks of the same cancer; 2) the same genes arenot necessarily implicated in familial clusters and sporadic cases (without a family history), even in the same population; and 3) familial cancers are relatively rare, and thus do not account for more than approximately 25% of all cases in a population, or 20% of incident breast cancers [12]. For these reasons, much doubt has been expressed over the last decade that family studies had much future utility for resolving complex genotypes for diseases like breast cancer[13]. Instead, as genome-wide data rapidly became available, and with it an acute need for “high-throughput” analyses, the focus of research quickly shifted to simpler association study designs to measure genetic differences between phenotypic classes, such as cases and controls.

The genome-wide association study (GWAS) approach focuses on genotype-phenotype co-variation, usually for a densely distributed set of SNPs over the genome. Positive associations occur where genotype differences correspond to phenotype differences outside of what is expected under a null hypothesis, and their locations mark points in or near genetic variants that cause disease or contribute to its risk. Numerous GWAS have been done in search of genes that condition risk of breast cancer, and a list of genes and variants with modest effects on cancer risk has certainly developed as a result [14] [15]. However, the small fraction of breast cancers attributable to these relatively common but low-penetrance alleles suggests that a larger set of genetic factors, more of them reaching moderate effect, but occurring with low frequency in a population, might account for such common cancer phenotypes. This “heritability gap” has been considered a problem of statistical lack of power to resolve a potentially large number of genetic variants, some of them low in frequency (rare), and of only moderate or low risk effect for common but deadly disease phenotypes.

For complex diseases in general, GWAS have generated many significant associations between particular SNPs and disease phenotypes, but again, these are often inconsistent across studies, populations, designs, and samples. After more than a decade of modeling and measuring complex genotype-phenotype associations by GWAS, it remains difficult to value the contributed effects of particular genes to a disease phenotype by this method, and today it is widely appreciated that the approach has a critical shortcoming. For many individual studies the methodology is simply underpowered to sort high throughput data for a definitive set of genetic factors—unknown in number and varying in frequency and effect size–responsible for a complex disease phenotype. As a result, there is much uncertainty about what an association study really captures, and clarification is often sought by improving power and reliability—by increasing genome coverage, sample size, or by meta-analyses. In this regard, today’s study designs are ambitious, involving huge numbers of cases and increasingly narrow definitions of the phenotype[16]. Even so, GWAS of breast cancer have not resolved single genes of major effect comparable to *BRCA1* and *BRCA2;* neither have they established a comprehensive predisposing genotype for the disease.

Although it is now considerably easier and less expensive to collect genetic data for GWAS, it has remained elusive by association testing to capture enough genetic variants, or of sufficient effect, to account for what is manifestly familial and estimated as heritable. In this study we address the notion of “missing heritability” and compromised analytic power for detecting genetic factors contributive to breast cancer. To do this, we have fashioned a “high-definition” approach to linkage analysis using deep pedigree data, albeit sparsely genotyped, and for pairs related over a range of relationships. The approach is not designed primarily to address the matter of heritability; more importantly, it is designed to advance the train of evidence leading to the identification of genetic variants that are potentially rare— i.e., found at low population frequency—of moderate effect on risk, and likely larger in number than the class of single genes of major effect, such as *BRCA1*.

## Subjects and Methods

### Study Sample: breast cancer cases from high risk families in Utah

The Utah Population Database (UPDB) is a repository of longitudinal information originally constructed from genealogical data pertaining to Utahans and their families [17]. Through successive record linking efforts, the database integrates cancer registry data, medical records data, Utah State certified deaths and births, etc. Currently, the UPDBcaptures information for approximately 7 million individuals, many of whom are members of extensive pedigree networks 2 to 14 generations deep [18]. Pedigree information from the UPDB, and Utah’s SEER cancer registry, were used to establish diagnosed breast cancer cases clustered in large multigenerational families. We then compared observed and expected breast cancer incidence in case families and recruited study subjects from “high risk” families, i.e., those with excess incidence having a probability of less than 0.01 of occurring by chance [19]. However, we excluded cases and families previously studied and known to be segregating *BRCA1* or *BRCA2* mutations as their primary genetic risk factors for breast cancer.

Female members of high risk families who were diagnosed with breast cancer and alive at the start of the study were invited to join, as were unaffected women drawn from the same large families. Study participants were home visited, at which time individual and family health histories were documented and blood samples collected (by venous puncture) as the source of DNA for genome-wide SNP analyses.

The genotyped study sample consisted of 154 women diagnosed with breast cancer, and 94 unaffected relatives. “Families” were defined after recruitment as the largest set of genotyped subjects, including a minimum of 3 cases, all descended from a common ancestor. By this method all participants (n=248) are members of 22 large families with evident excess risk of breast cancer. Cases (n=154) collectively form 1,011 affected relative (AA) pairs for linkage analysis; genotypes of unaffected subjects (94) were used to estimate allele frequencies and identity by descent probabilities. The families included in this study are pictured schematically in Figure 1.

**Figure 1.**
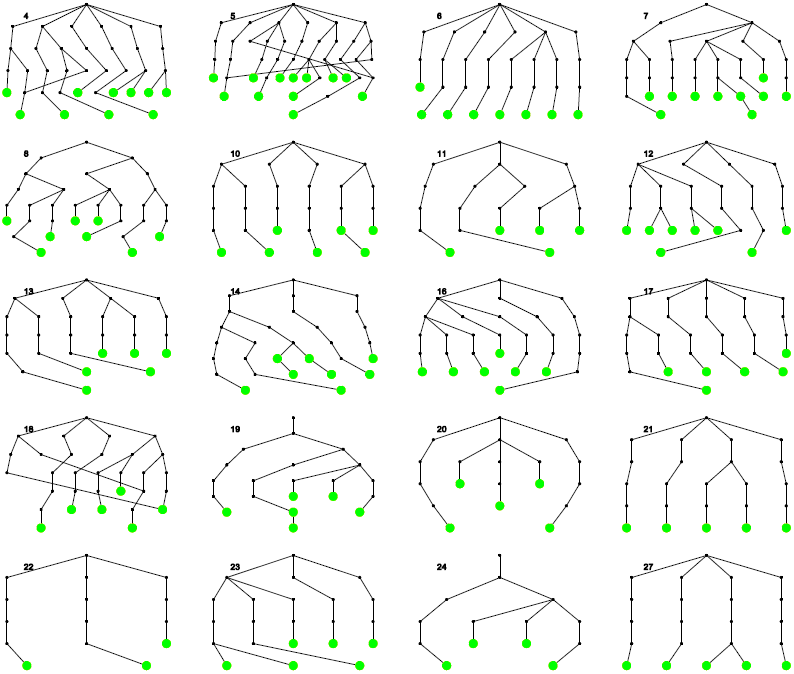
Schematic pedigrees of the 22 families studied. Affected subjects are indicated with enlarged green dots. Only lines of descent from common ancestors are shown. Crossing lines indicate inbreeding, although only one affected subject was herself inbred (family 18). Family numbers 2, 9, 15, 25 and 26 were assigned to families not used for this analysis, either because of overlap with another family (2), or insufficient number of usable samples from breast cancer cases.

The University of Utah Health Sciences Institutional Review Board and the University of Louisville Biomedical Institutional Review Board approved the study protocol; all recruited subjects provided their written consent to be included in this study.

### Genotypes

Genotyping was performed with Illumina 370 Duo and 610 Quad arrays at deCODE Genetics, Reykjavik, Iceland. SNPs with low quality scores (GenCall[20] quality score <0.15), and those with inconsistent allele frequencies between the two arrays (any absolute difference in minor allele frequency > 0.05), were eliminated. All alleles were called on the forward strand, and checked for consistency between arrays. After approximately 15% of the SNPs were eliminated by these quality control criteria, a total of 285,630 genotypes per subject were retained. Mendelian consistency checks were not performed because of the very small number of families with informative data.

### Evaluation of genetic vs. genealogical relatedness

We examined the degree to which relatedness assessed by genome-wide genetic similarity corresponded to relatedness as reported in the UPDB genealogical data for pairs of relatives. For this study, we used genotypes on 429 individuals, including the 248 subjects in the linkage study, as well as 181 women from families with fewer than 3 genotyped breast cancer cases. A total of 91,806 pairs were evaluated, using coefficient of relatedness to characterize the genealogical data, and the genetic relatedness matrix computed by GCTA[21] to characterize relatedness from SNP data. To facilitate comparison, relatedness from each measure was grouped by rounding −log_2_(relatedness) to correspond to degree of relationship.

### Identity by Descent (IBD) estimation for linkage analysis

Pairs formed from the sample set were used to generate Identity by Descent (IBD) matrices for linkage analysis. IBD was computed using PEDIBD software developed by Li and colleagues[22]. Their method employs a Viterbi algorithm [23] to find the most likely path of descent of an ancestral allele through a deep, but sparsely genotyped pedigree structure, via hidden Markov models of inheritance and recombination. The method efficiently parses the high-density genotype data of the Illumina arrays, permitting estimation of IBD matrices for 1,011 affected relative pairs at up to 285,630 loci in approximately 24 hours of CPU time on current equipment (substantially less for thinned data sets). Allele frequencies were estimated by simple counting among all genotyped individuals, affected or unaffected. As noted by Boehnke[24] and others, simple counting among family members does not introduce any systematic bias in the absence of allelic association, and any association would introduce a conservative bias as it would lead to overestimation of the frequency of a disease-associated allele.

### Test statistic for linkage

We employed the IBDREG quasi-likelihood approach described by Schaid, et al. [25] to test for concordant pair (affected only) linkage without covariates. IBDREG has an important advantage in comparison to competing methods as it appropriately adjusts for between-pair covariance when multiple relative pairs are drawn from the same pedigree structure. Because the families studied vary considerably in size, and some have only a few affected members, the distributional properties (and hence the asymptotic p-values) of the test statistic were uncertain. Therefore, we used simulation to compute p-values and family-wise error rates. The approach is described below.

### Simulation of Identity by Descent in the Absence of Linkage, but the Presence of Linkage Disequilibrium

We performed 100 full-genome simulations of identity by descent using all 285,630 autosomal markers and all 22 families for three reasons: 1) to allow accurate estimation of error rates for IBD estimates across all family structures and all autosomes; 2) to give a reference against which different thinning strategies could be evaluated for their effects on both IBD accuracy and the distribution of the linkage test statistic; and 3) to provide distributions of the test statistic under the null hypothesis.

### Estimation of error rates

It is well known that linkage analysis based on high-density SNP arrays is subject to potentially severe bias away from the null because of linkage disequilibrium (LD). LD between nearby markers will cause overestimation of the probability that two related individuals share marker alleles that are identical by descent (IBD) [26, 27]. Although in principle, simultaneous modeling of LD among founders and IBD among descendants would be the most powerful approach to using all the genotype data at our disposal [28], the computational burden of such modeling in complex multigenerational families is not readily surmountable at present.

### Marker thinning intervals

Marker thinning effectively varies the strength of LD by setting maximum R^2^ between SNPs at various thresholds (0.6, 0.4, and0.2 here). At each threshold, SNPs were thinned by recursively finding the midpoint of a block of SNPs mutually correlated at R^2^ > the current threshold, then dropping all but the midpoint SNP, so that the maximum pairwise correlation could not exceed the selected level. Thinned marker sets were run against simulated (null) genotype data for chromosome 7 to establish error rates in the IBD estimates and thus, the contribution to false positive linkage scores for varying strengths of LD structure.

For simulation analyses, we imputed an LD structure descending from founders by adapting the HapMap3, Phase 2 observed LD structure for 234 independent haplotypes estimated from 117 CEU subjects [29]. The HapMap sample series is appropriate as a reference set for this study because it too is a Utah family series [30]. HAPGEN2 software [31] was used to generate 4000 random haplotypes with the desired LD characteristics for all 285,630 autosomal loci. For each pedigree founder, two random haplotypes were chosen, from the 4000 randomly generated, by sampling with replacement. Alleles for each SNP marker were randomly generated in proportion to each marker’s allele frequencies. Haplotypes were descended through the study pedigrees, resetting random segregation indicators according to HapMap’s estimated recombination fractions. Recombination between markers was estimated by cubic spline interpolation using R [32].

Pedigree information and simulated marker data were input to PEDIBD to obtain a full matrix of IBD estimates for all affected pairs. IBD estimates generated by PEDIBD were compared to the simulated “true” IBD states (0, 1, or 2 alleles known to be shared for each pair) to determine error rates.

### Distribution of test statistics under the null

The IBD estimates generated by PEDIBD were input to IBDREG to calculate linkage statistics. All simulated IBD states and marker allele data were generated under the null hypothesis of no linkage between marker loci and disease predisposition. Thus, the distribution of test statistics for each marker locus within each family can be taken to represent a sample from the null distribution for a whole genome scan of that family. In addition to the per-locus asymptotic p-value computed by IBDREG, we report a family-specific per-locus Monte Carlo p-value, a family-specific per-genome Monte Carlo p-value, and a Monte Carlo composite false discovery rate (FDR) controlling for the whole-genome analysis of 22 families[33].

### Identification of linkage peaks and boundaries

We defined a putative linkage peak as the chromosomal location of the smallest p-value over a run of consecutive SNPs with asymptotic p-values less than 0.001. The extent of the linked “peak” region was identified from the focal SNP (smallest p-value) to the nearest SNP either side with a p-value tenfold greater than the focal SNP, thus establishing the boundary maximum *p*. Overlapping peaks across multiple families were counted as a single peak.

## Results

An initial check for correspondence between coefficients of relatedness estimated from pedigree information and from SNP genotypes was made for all possible pairs of study subjects (see Methods). This information is plotted in Figure 2 for pairs of related individuals. The most distantly related pairs in the genealogical data were 13^th^ degree relatives, so pairs unrelated by genealogy and pairs estimated to be genetically more distant than 13^th^ degree were plotted as though they were 14^th^ degree relatives on either scale. There was generally very good agreement between genealogical and genetic distance up to about the 6^th^ degree, and a gradual loss of precision past that point in this population.

**Figure 2.**
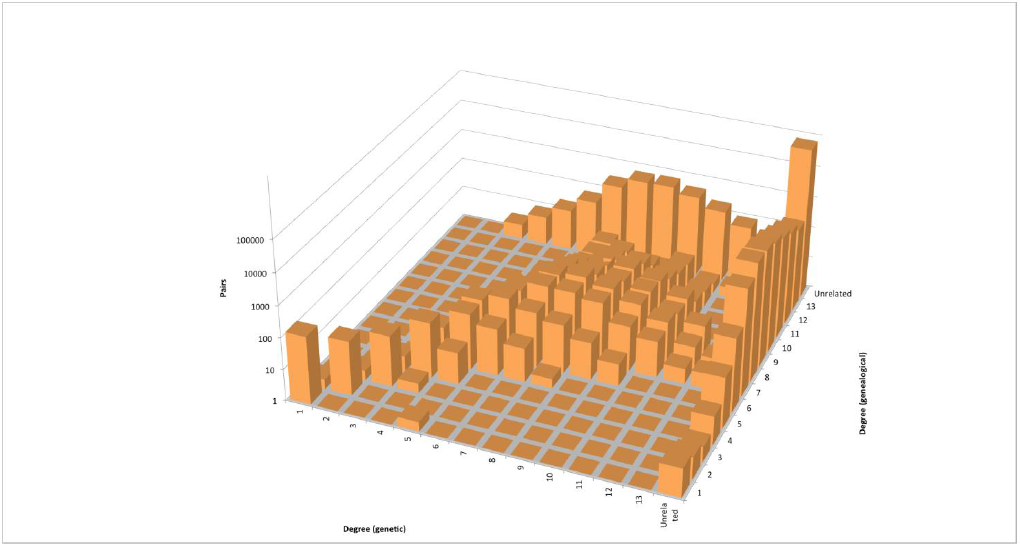
Genetic vs. genealogical relatedness. Relatedness estimated as global IBD from genetic data (SNPs) compared to genealogical relatedness (from pedigrees) for all possible pairs of study subjects (affected and unaffected). Red dots indicate pairs with substantial mismatch between genealogical and genetic distances; these pairs were dropped from the analysis by inspection and removal of one or both subjects from pedigree data.

It is common that some members of large Utah families overlap in family membership in descending generations, and Table 1 gives counts of these individuals. Note that most subjects are members of only one family, and the majority of those who overlap in family membership do so as pedigree members, rather than genotyped study subjects. This is shown in Table 1. where counts are given for the number of individuals with membership in >1 of the 22 families, by disease status. Counts of individuals and affected pairs by family are given in Table 2.

**Table 1.**
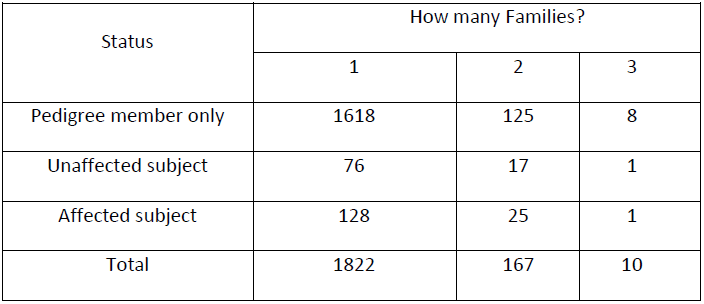
Number of individuals with membership in ≥1 of the 22 family groups, by disease status.

| Status | How many Families? |  |  |
| --- | --- | --- | --- |
|  | 1 | 2 | 3 |
| Pedigree member only | 1618 | 125 | 8 |
| Unaffected subject | 76 | 17 | 1 |
| Affected subject | 128 | 25 | 1 |
| Total | 1822 | 167 | 10 |

**Table 2.** Total number of affected study subjects per family, and number of pairs per family for linkage analysis.

| Family<sup>a</sup> | Affected<br>Individuals | Pairs |
| --- | --- | --- |
| 1 | 31 | 465 |
| 3 | 16 | 120 |
| 4 | 11 | 55 |
| 5 | 11 | 55 |
| 6 | 8 | 28 |
| 7 | 10 | 45 |
| 8 | 8 | 28 |
| 10 | 7 | 21 |
| 11 | 5 | 10 |
| 12 | 8 | 28 |
| 13 | 6 | 15 |
| 14 | 7 | 21 |
| 16 | 7 | 21 |
| 17 | 6 | 15 |
| 18 | 6 | 15 |
| 19 | 6 | 15 |
| 20 | 5 | 10 |
| 21 | 5 | 10 |
| 22 | 3 | 3 |
| 23 | 6 | 15 |
| 24 | 4 | 6 |
| 27 | 5 | 10 |
| Total | 181 | 1011 |
| Count<sup>b</sup> | 154 |  |

<sup>a</sup>Families are numbered to 27, but 2, 9, 15, 25, and 26 were not included in the study; total families = 22.
<sup>b</sup>The total number of distinct individuals. Some subjects were members of more than one family (see Table 1).

Simulations were done to depict the inflationary effect of LD on IBD and false positive linkage scores (see **Methods**). These results are shown in Figure 3 and Table 3. In order to control for this effect, and reduce false positive linkage signal, SNPs were thinned to various thresholds of correlation between them. At the threshold R^2^ ≤ 0.4, IBD over-estimation due to LD was controlled fairly well, but positive linkage peaks still occurred. At R^2^≤ 0.2, spurious linkage peaks disappeared. The results given in Figure 4 and Table 4 are based on the inter-marker threshold R^2^ ≤ 0.2 for the thinned set of 19,609 SNPs.

**Figure 3.**
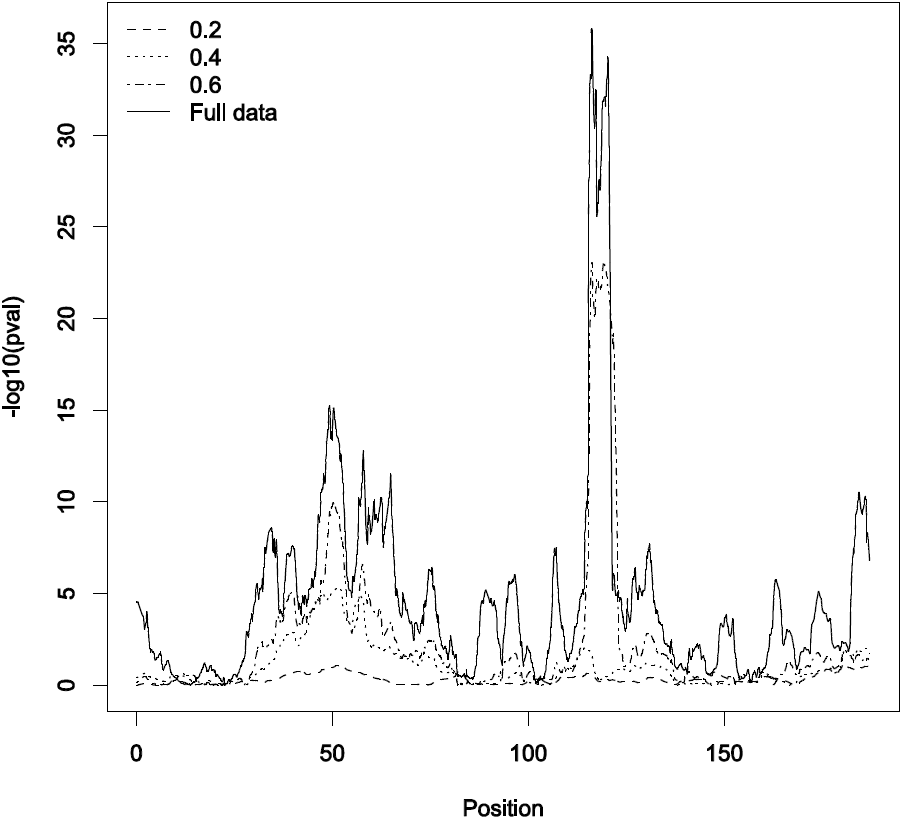
Null simulation results for chromosome 7 markers. False positive linkage peaks from simulation of null linkage at varying LD thinning thresholds on chromosome 7.

**Figure 4.**
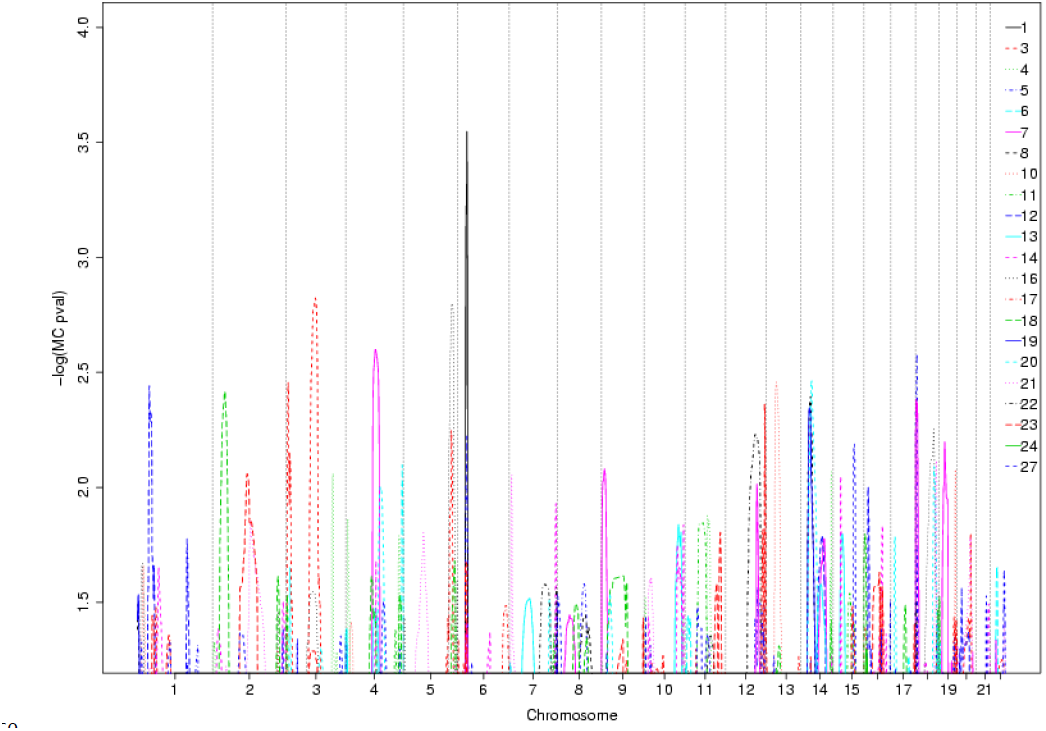
Linkage peaks by chromosome and family. Only unadjusted p-values < 0.1 are displayed. Family numbers (legend) correspond to those shown in Figure 1.

**Table 3.**
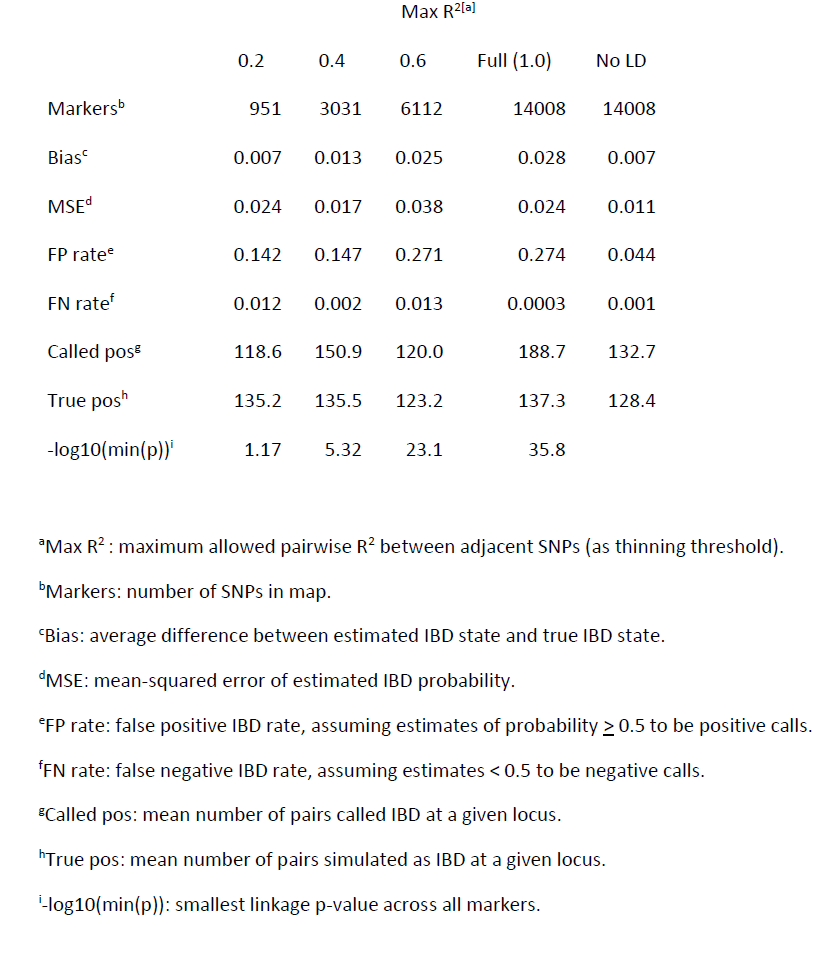
Summary of null simulation results for chromosome 7 at various thinning intervals.

| | Max $R^{2[a]}$ | | | | |
| --- | --- | --- | --- | --- | --- |
|  | 0.2 | 0.4 | 0.6 | Full (1.0) | No LD |
| Markers<sup>b</sup> | 951 | 3031 | 6112 | 14008 | 14008 |
| Bias<sup>c</sup> | 0.007 | 0.013 | 0.025 | 0.028 | 0.007 |
| MSE<sup>d</sup> | 0.024 | 0.017 | 0.038 | 0.024 | 0.011 |
| FP rate<sup>e</sup> | 0.142 | 0.147 | 0.271 | 0.274 | 0.044 |
| FN rate<sup>f</sup> | 0.012 | 0.002 | 0.013 | 0.0003 | 0.001 |
| Called pos<sup>g</sup> | 118.6 | 150.9 | 120.0 | 188.7 | 132.7 |
| True pos<sup>h</sup> | 135.2 | 135.5 | 123.2 | 137.3 | 128.4 |
| $-\log_{10}(\min(p))$<sup>i</sup> | 1.17 | 5.32 | 23.1 | 35.8 | |

<sup>a</sup>Max $R^2$ : maximum allowed pairwise $R^2$ between adjacent SNPs (as thinning threshold).
<sup>b</sup>Markers: number of SNPs in map.
<sup>c</sup>Bias: average difference between estimated IBD state and true IBD state.
<sup>d</sup>MSE: mean-squared error of estimated IBD probability.
<sup>e</sup>FP rate: false positive IBD rate, assuming estimates of probability $\geq 0.5$ to be positive calls.
<sup>f</sup>FN rate: false negative IBD rate, assuming estimates $< 0.5$ to be negative calls.
<sup>g</sup>Called pos: mean number of pairs called IBD at a given locus.
<sup>h</sup>True pos: mean number of pairs simulated as IBD at a given locus.
<sup>i</sup> $-\log_{10}(\min(p))$ : smallest linkage p-value across all markers.

**Table 4.**
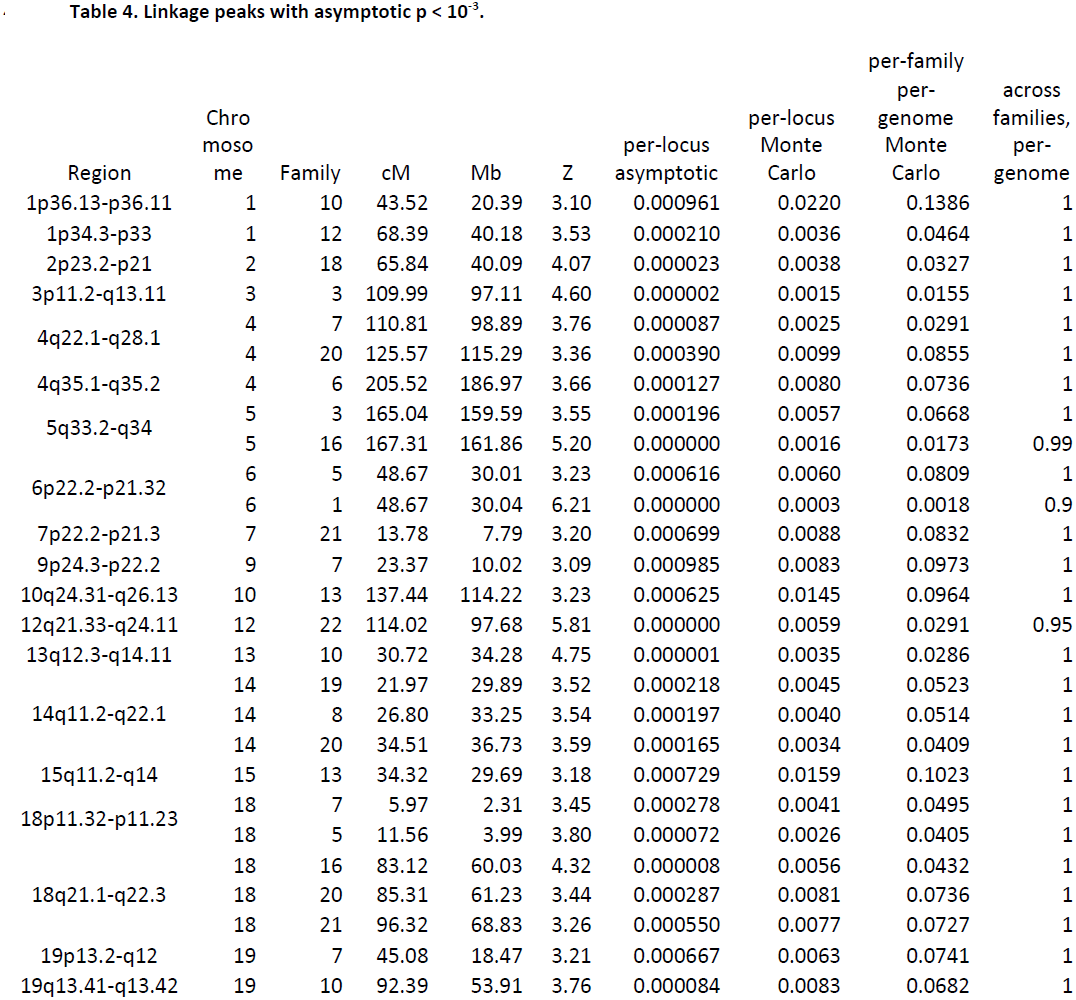
Linkage peaks with asymptotic p < 10^−3^.

Table 4 gives linkage results for 1,011 affected relative pairs generated from a total of 154 genotyped breast cancer cases. The analysis identified 19 distinct peaks with asymptotic unadjusted within-family p < 0.001. More realistic estimates of the probability of these results under the null hypothesis are derived from the 100 per-family genome-wide simulations, and presented in Table 4 as well. Monte Carlo per-locus p-values are generally considerably larger than the asymptotic p-values, particularly for smaller families. After further adjustment for genome-wide comparisons within families, 11 regions retained adjusted p-values below 0.05, and 17 regions retained adjusted p-values below 0.1. However, when we adjusted for simultaneous whole-genome search across all 22 families, only the 3 peaks with the highest scores were large enough that a single random result under the null would not have been expected to exceed them 100% of the time.

**Supplementary Table 1** lists all breast cancer-associated genes from DisGeNET

[http://www.disgenet.org/web/DisGeNET/v2.1], TCGA [34] and Cancer Resource[35] located within peaks defined by a 10-fold increase in asymptotic p-value. The large peak on chromosome 6 for family 1 includes multiple genes that have been associated with breast cancer risk and/or tumorigenesis, including members of the *HLA* complex, *NOTCH4,* and *TNF,* among others. Also noteworthy is that the chromosome 13 peak for family 10 includes *BRCA2,* while no family exhibited linkage to the *TP53* or *BRCA1* regions on chromosome 17. Figure 4 shows the relative locations and amplitudes of the linkage peaks by family.

## Discussion

It is low-frequency variants that are difficult to find in convincing association with a disease phenotype from genome-wide association tests[13]. However, if we are to resolve this low frequency, moderate risk class of variants, then population-wide sampling from whole undifferentiated, or minimally structured populations, is perhaps not the most strategic sampling approach to use. Variants of this class occur de novo, are replicated and transmitted to descendants. For this reason, they will reach their highest frequencies within family lineages[36], the larger the better, while remaining at low frequency (rare) in any usual population sample, whether n = 100s or 100,000s. The moderate risk nature of this class of variants is likely due to the fact that their risk effects depend on participation in larger gene networks to account for increased cancer risk in particular families. In this sense, variants of smaller effect can alter disease risk in the context of gene networks that regulate the functional pathways involved in the onset and/or progression of the disease.

The family study approach does not rest on anticipating “a new breast cancer genotype”, nor a “comprehensive genotype” to account for breast cancer risk in this population and by the usual purview of linkage analysis. Instead, we tried to capture evidence of low frequency variants at the population level, but enriched at the level of very large high-risk families. Our approach yielded 17 genomic regions possibly linked (per-family per-genome Monte Carlo p ≤ 0.1) to breast cancer for the 22 families studied, but with considerable variation among families: 15 of the 22 families (68%) showed possible linkage to at least 1 region by the criteria used here; 1 family showed possible linkage to 4 regions; 1 family to 3 regions; 5 families to 2 regions; and 8 families showed linkage to 1 region. It is noteworthy that linkage to the *BRCA2* region on chromosome 13 was observed in only one family (10), while no family exhibited linkage to the *BRCA1* region on chromosome 17.

The availability of high-density marker sets, efficient algorithms for estimating IBD in large families, and substantial computational resources permitted simulation of 100 null genome-wide results for each family. The simulation results then allowed us to compare Monte Carlo p-values with asymptotic p-values based on large sample theory. In Table 4 it is shown that the asymptotic estimates are frequently smaller than the Monte Carlo p-values by an order of magnitude or more. The genome-wide results for each family represent an appropriate basis for comparison to other published results based on linkage studies of one or a few families. Adjusting linkage estimates for all 22 families simultaneously, we find no linkage scores, or peaks, that could not have occurred by chance: the observed Z-score of 6.21 for family 1 on 6p21-22 was exceeded in 90% of null simulations—for at least one family at some location over the genome. However, in the simulated data, anomalously high Z-scores were much more commonly observed in small families—and never in family 1—so the adjustment across families is in some part size dependent, and therefore, less than perfect. For this reason, it might be more appropriate to consider the simulated probability of observing the result in 22 families just like family 1, which would be approximately 1-(1-0.0018)^^^22 = 0.039. Moreover, the existence of a possible linkage peak for family 5 in precisely the same location as family 1 on 6p21-22 strengthens the case for a susceptibility locus in this region.

For heuristic purposes, we can combine multiple lines of evidence to rank the various linked regions by priority. First, regions that overlap across multiple families (e.g. 6p22-21, families 5 and 1; 18p11, families 5 and 16; 18q21-22, families 20 and 21) likely indicate either a shared disease-predisposing haplotype inherited from an unknown common ancestor, or multiple predisposing variants in the same gene in truly unrelated, or variably related, families. Next, per-family, per-genome Monte Carlo p-values well below 0.05 (e.g. 3p11-13, family 3; 12q21-24, family 22) warrant further investigation. Finally, linked regions in individual families (13q12-14, family 10) that overlap with known breast cancer-predisposing loci, i.e. BRCA2, have the potential to greatly simplify mapping variants associated with specific diseases. In addition, the other regions suggestive of linkage without known breast cancer associated variants, might provide useful new clues about the location of genetic variants that increase the risk of breast cancer in members of these families, and serve as evidence of residual heterogeneity in genomic regions responsible for familial cancer susceptibility.

Although this linkage analysis was meant to identify regions of the genome that include putative genetic contributors to disease, there is still considerable distance between regions identified by linkage, and discovery of whether or what variants within them contribute to breast cancer risk (“true positives”). Having identified segments of the genome smaller than the whole, there are still at least six to ten segments to consider, each spanning many genes, a large amount of information and a lot of variation. Depending upon the definition of peak region—whether 1Mb-5Mb surrounding the focal SNP, or the larger regions bounded by a tenfold change in p-value—many genes that have been associated with breast cancer risk in other studies are captured in the linkage regions (Table S1), including BRCA2. From functional annotation, the linked regions we have identified encompass many genes that “look good” as candidates for further analysis. However, in order to identify specific variants relevant to breast cancer phenotypes in this study, especially those that are rare and of obscure overall effect, it remains to further interrogate the linkage regions by sequencing. An efficient approach might begin with whole exome sequencing to address functional variants first. Regional, functional, family and pair-specific information can all be used to direct targeted evaluations of concordance between expected linkage (SNP-based probabilities of sharing IBD) generated by our model, and differences in sequence sharing per exome through the linked regions. By using the linkage-partitioned information thus far, sequencing should reveal more specifically the locations of rarer variants likely relevant to the disease. Linkage and sequencing techniques together should do much to clarify the genetic architecture of breast cancer in this population, its heterogeneity among families[37], and importantly, give us a deeper understanding of the role of rare variants in conditioning risk differently among groups.

For any novel variants that might be established, or candidate genes that might be confirmed with sequencing, the hope is that the information will advance knowledge of the genetic pathways involved and their interacting factors so essential to personalized therapies, management, and outcomes in the clinical setting. The developing field of Molecular Epidemiology and its unique integrative approach to medical research only begins with the address of large and growing quantities of data for translation to improved risk prediction. Studies of this type, that inform us about what specific genomic variation underlies risk variation in a population, lead to the identification of risk subgroups, and most importantly, high-risk families and individuals. New and abundant genetic information will no doubt lead us to understand important features of how genes—common, rare, and in multiplicity—contribute to disease spectra, from well to mortal, and the intermediate.

## Supplemental Table and Figures

**Figure S1.**
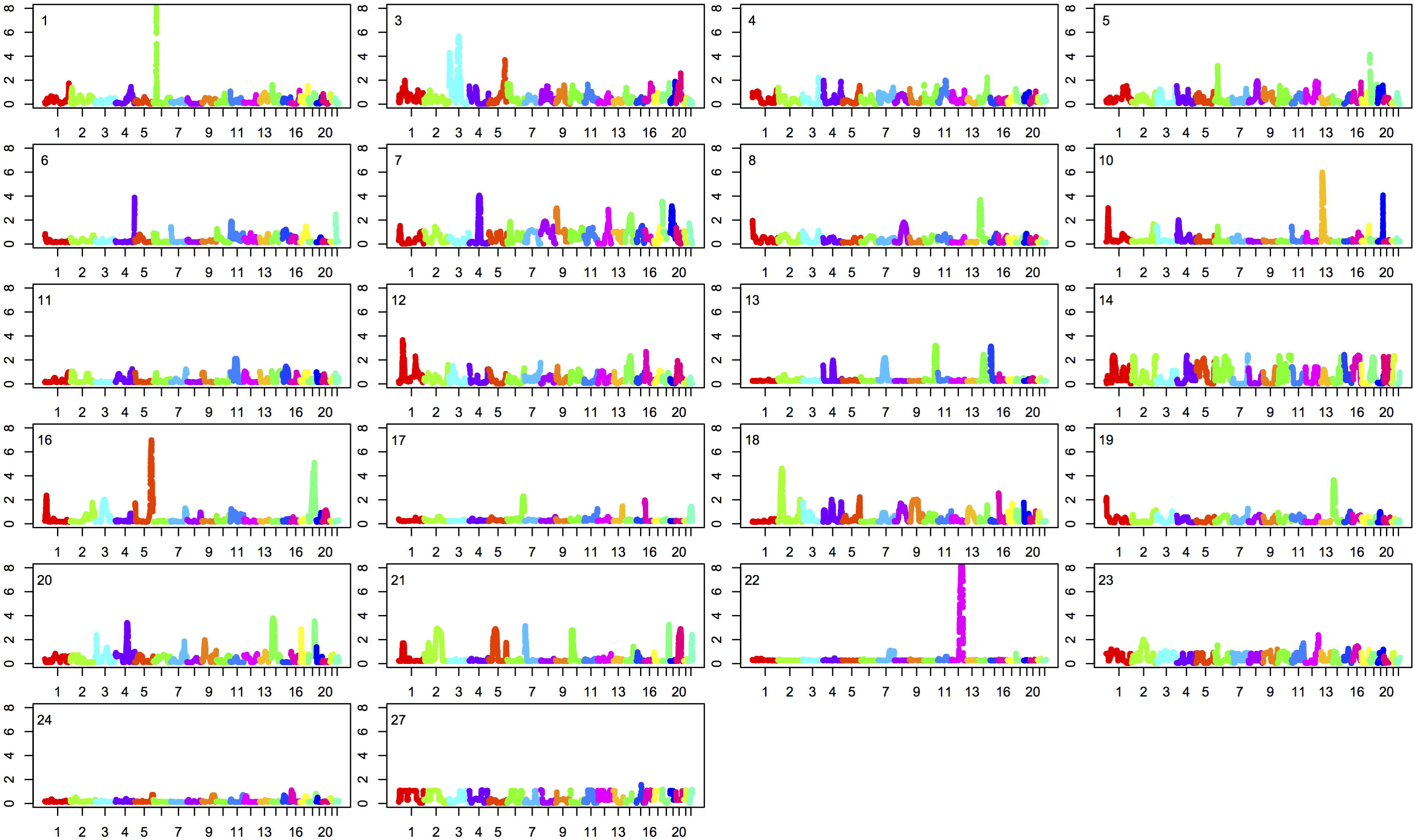
Manhattan plots for each family. Families are labeled as per Figure 1 in upper right corner of each plot.

**Figure S2.**
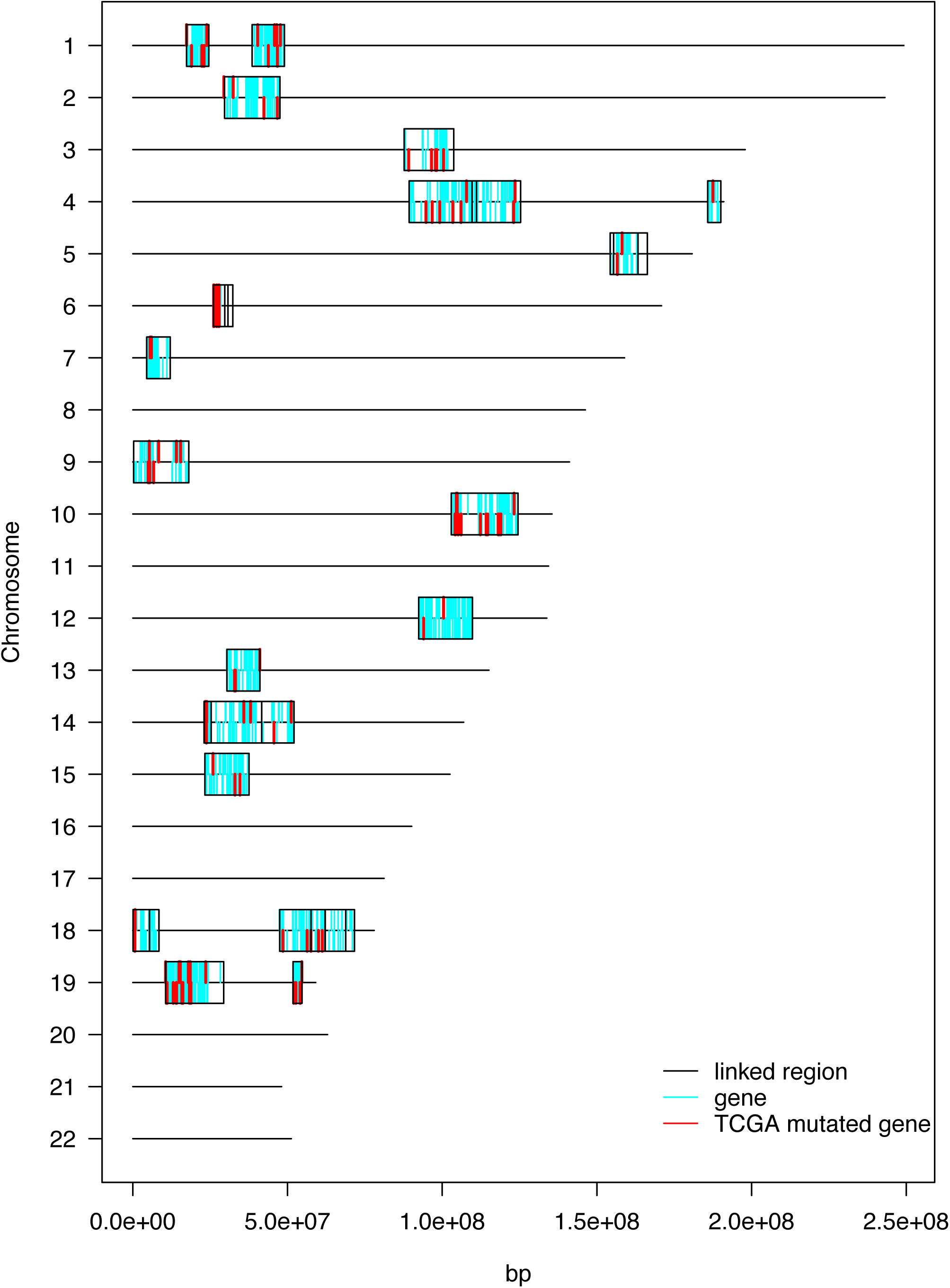
Locations of genes within linkage peaks with unadjusted p-value < 0.001. Within peaks, cyan lines indicate genes, red lines indicate genes mutated in TCGA breast cancer specimens, black lines indicate boundaries of overlapping peaks. Coding strand is indicated by placement within box: genes coded on the forward strand are drawn above the midline, while genes coded on the reverse strand are drawn below.

Table S1. Table of all genes in linked regions, ordered by bioinformatics resource scores (see text for references): tcga.mut = number of mutations observed in TCGA breast tumors; cr = cancer resource breast cancer associated (1) vs not associated (0); dg = disGeNet breast cancer association score; sum = sum((tcga.mut > 0)+(cr > 0)+(dg > 0)); tcga = TCGA breast tumor mutations/bp.

## Acknowledgments.

Data analysis and genotyping was supported by Susan G. Komen Foundation grant KG-071514. Partial support for all datasets within the Utah Population Database was provided by the University of Utah Huntsman Cancer Institute and the Huntsman Cancer Institute Cancer Center Support grant, P30-CA2014 from the National Cancer Institute. Support for the Utah Cancer Registry is provided by Contract No HHSN2612013000171 from the National Cancer Institute with additional support from the Utah Department of Health. We thank Debbie Shea, Matt Hoffman, Terah Young, and Loren Budge for subject recruiting and data collection; Geri Mineau, Alison Fraser and Janice Conrads at the Huntsman Cancer Institute’s Pedigree and Population Resource for linking subject data to the Utah Population Database; Xin Li for consultation on PEDIBD modifications for application to our data; Dan Schaid, Duncan Thomas, Steve Schrodi, and Ken Boucher for valuable discussions of this work.

